# Optimal dimensionality selection for independent component analysis of transcriptomic data

**DOI:** 10.1101/2021.05.26.445885

**Authors:** John Luke McConn, Cameron R. Lamoureux, Saugat Poudel, Bernhard O. Palsson, Anand V. Sastry

## Abstract

Independent Component Analysis (ICA) is an unsupervised machine learning algorithm that separates a set of mixed signals into a set of statistically independent source signals. Applied to high-quality gene expression datasets, ICA effectively reveals the source signals of the transcriptome as groups of co-regulated genes and their corresponding activities across diverse growth conditions. Two major variables that affect the output of ICA are the diversity and scope of the underlying data, and the user-defined number of independent components, or dimensionality, to compute. Availability of high-quality transcriptomic datasets has grown exponentially as high-throughput technologies have advanced; however, optimal dimensionality selection remains an open question. Here, we introduce a new method, called OptICA, for effectively finding the optimal dimensionality that consistently maximizes the number of biologically relevant components revealed while minimizing the potential for over-decomposition. We show that OptICA outperforms two previously proposed methods for selecting the number of independent components across four transcriptomic databases of varying sizes. OptICA avoids both over-decomposition and under-decomposition of transcriptomic datasets resulting in the best representation of the organism’s underlying transcriptional regulatory network.

## Introduction

Independent Component Analysis (ICA) is an unsupervised machine learning algorithm which models a multivariate dataset as a linear combination of statistically independent hidden factors or components [1]. For example, ICA may be used to solve the cocktail party problem, in which multiple mixed audio signals (i.e., people speaking simultaneously at a cocktail party) are recorded by microphones dispersed throughout a room. Each device records a unique linear mixture of the original signals depending on its proximity to each speaker. Applying ICA to this set of mixed recordings can effectively recover the original independent audio signals, and their relative volumes for each microphone.

Beyond deconvoluting audio signals, ICA is widely applicable to several other fields involving signal separation or feature extraction [2, 3]. With the advancement of high-throughput gene expression profiling, ICA has proven to be useful in analyzing highly multivariate microarray and RNA sequencing (RNA-seq) gene expression datasets [4–8].

ICA has been applied to large microbial transcriptomics datasets, resulting in highly accurate reconstructions of their underlying transcriptional regulatory networks (TRNs). For example, ICA decomposition of an *Escherichia coli* expression compendium containing 278 expression profiles (named PRECISE) revealed 92 independently modulated gene sets, termed iModulons. Most iModulons significantly overlapped with known regulons and could be directly linked to a single transcription factor [4]. Similar analysis has since been carried out on a *Bacillus subtilis* microarray dataset with 269 expression profiles [9] and on a compendium of 108 RNA-seq profiles of *Staphylococcus aureus*, named *Staph*PRECISE [10], which revealed 83 and 29 similarly informative iModulons, respectively. Although independent components extracted from human datasets also capture biologically relevant gene clusters [11, 12], the similarity between these gene clusters and the known TRN is obscured by the inherent complexity of eukaryotic transcriptional regulation.

As gene expression datasets incorporate more individual growth conditions, it becomes possible to develop a comprehensive reconstruction of an organism’s TRN. To achieve this, the output of ICA decompositions depends primarily on two inputs – the number of high-quality data sources across diverse growth conditions; and the number of independent components to compute. While the former has become more accessible through public data repositories [13], determining the optimal dimensions for the decomposition remains an open question [14]. Previous investigation into this problem showed that searching for too many components can result in many components driven by small gene sets, whereas searching for too few components can obscure the biological interpretation [15].

Several methods have been suggested and employed to answer this question. One such method entails setting the number of dimensions equal to the number of principal components, determined through principal component analysis (PCA), which account for a certain level of variance in the data [4, 8]. Alternatively, the Maximally Stable Transcriptome Dimension (MSTD) has been suggested, defined as the maximum dimension before ICA begins to produce a large proportion of unstable components [15]. However, these methods have not yet been rigorously tested against well-characterized microbial TRNs.

In this study, we investigated how different dimensions for ICA affect the accuracy of the inferred TRN and evaluated the performance of existing dimensionality selection methods. We found that these previously proposed selection methods were inconsistent, resulting in either over-decomposition or under-decomposition in various transcriptomic datasets. From these results, we developed a new method that identifies the optimal dimension that maximizes independent components that represent known regulation, while minimizing the presence of biologically meaningless over-decomposed components. The new method, named OptICA, ensures that future studies will select the ideal dimensionality to optimize the reconstructed TRNs from new transcriptomic datasets.

## Results

### Independent components form a “tree” across dimensions

To develop an understanding of how independent components evolve across different dimensions, we decomposed four transcriptomic datasets (Table 1) using FastICA [16] across the full range of possible dimensions. Since FastICA is inherently stochastic, we applied a clustering approach that only retains independent components that persist across multiple runs (see Methods). Therefore, the number of robust components identified at a particular dimension may be lower than expected, as unstable components are discarded (**Figure S1**). Application of a threshold to each of these robust components resulted in discrete groups of genes for each component, named iModulons.

Several RNA-seq and microarray datasets were utilized for this analysis, including the original version of PRECISE (PRECISE 1.0) [4], an expanded version (PRECISE 2.0), a compendium of *S. aureus* RNA-seq expression profiles (StaphPRECISE) [10] and a *B. subtilis* microarray dataset [9, 17] (Table 1). Each dataset was decomposed from two dimensions through full decomposition (one dimension for each expression profile), and independent components were compared between adjacent dimensions to form a “dimensionality tree”.

Dimensionality trees convey the evolution of the independent component structure across dimensions, as shown by the dimensionality tree for PRECISE 1.0 (**Figure 1**). At low dimensions, only a few independent components are identified. The iModulons derived from these independent components often contain multiple related regulons (**Figure 1a**). As the number of dimensions increases, these iModulons tend to split such that each regulon is contained within its own iModulon. As existing components split and new components appear, there is a net increase in total independent components until a relatively stable decomposition structure is reached. The appearance of additional components beyond this stable region suggests the commencement of over-decomposition evidenced by the appearance of components with a single highly weighted gene (**Figure 1b**).

**Figure 1.**
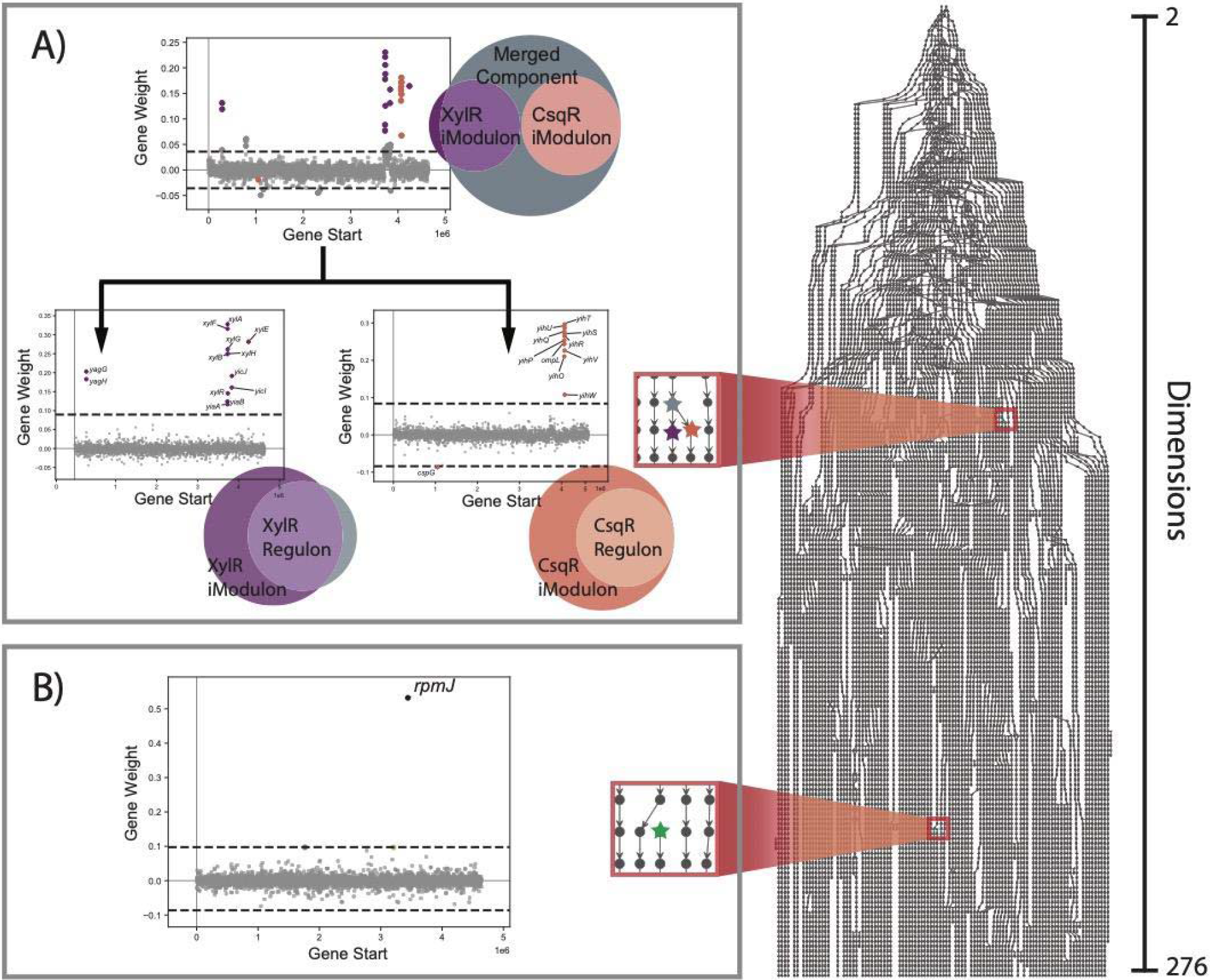
The dimensionality tree of PRECISE 1.0 reveals how its ICA decomposition evolves across a range of dimensions. Each point represents a computed component, and a row of points represents all components calculated at a particular dimensionality. Connections between components of adjacent dimensions were established where their correlation was greater than 0.3. **(A)** At low dimensions, components undergo splitting whereby disparate gene sets, initially contained in a single component, split into multiple components which more accurately reflect underlying transcriptional regulatory mechanisms. **(B)** At high dimensions new components appear which are uncorrelated to those of the preceding dimension. Often, these components contain a single highly weighted gene and signify the commencement of over decomposition.

Dimensionality trees were computed for all four transcriptomic datasets and demonstrated how the overall structure of the ICA decomposition evolves as more components are computed. With few exceptions, once a component was initially discovered at a particular dimension, it proved to be conserved in higher order decompositions (**Figure S2**). This realization suggests that across the stable decomposition region, the independent component structure does not materially change. Dimensionality selection techniques that produce useful decompositions of transcriptomic data would likely target points within this range, following the initial linear increase in computed components but before the commencement of over decomposition.

### Dimension selection methods often result in over or under decomposition

To identify the optimal dimension, the information from the dimensionality tree was summarized into various categories of iModulons. An iModulon was classified as “regulatory” if it was significantly enriched with a specific regulon (Fisher’s exact test, FDR < 1e-5). Since iModulons found at high dimensions are often highly similar to known regulons, we also tracked the number of “conserved” iModulons, or iModulons at lower dimensions that were similar to an iModulon detected at the largest dimension. Finally, to capture over-decomposition, we tracked the number of iModulons that contained a single highly weighted gene, and conversely, contained more than one gene. The categories were not exclusive; an iModulon at a particular dimension could be a regulatory, non-single gene, conserved iModulon.

These classifications were plotted across the full range of possible dimensionalities for the four datasets (**Figure 2**). These charts clearly showed that the number of regulatory iModulons and non-single gene iModulons sharply increased at lower dimensions. For the larger datasets (PRECISE 1.0 and 2.0), the number of regulatory iModulons and non-single gene iModulons plateaued and did not substantially increase even at the full decomposition. However, the number of conserved iModulons increased during plateau, indicating that the iModulons were experiencing internal reorganization. On the other hand, we observed large numbers of single-gene iModulons at high dimensions, signifying over-decomposition. For PRECISE 2.0, the largest dataset in this study, nearly all new components above a dimension of 500 were single-gene components.

**Figure 2.**
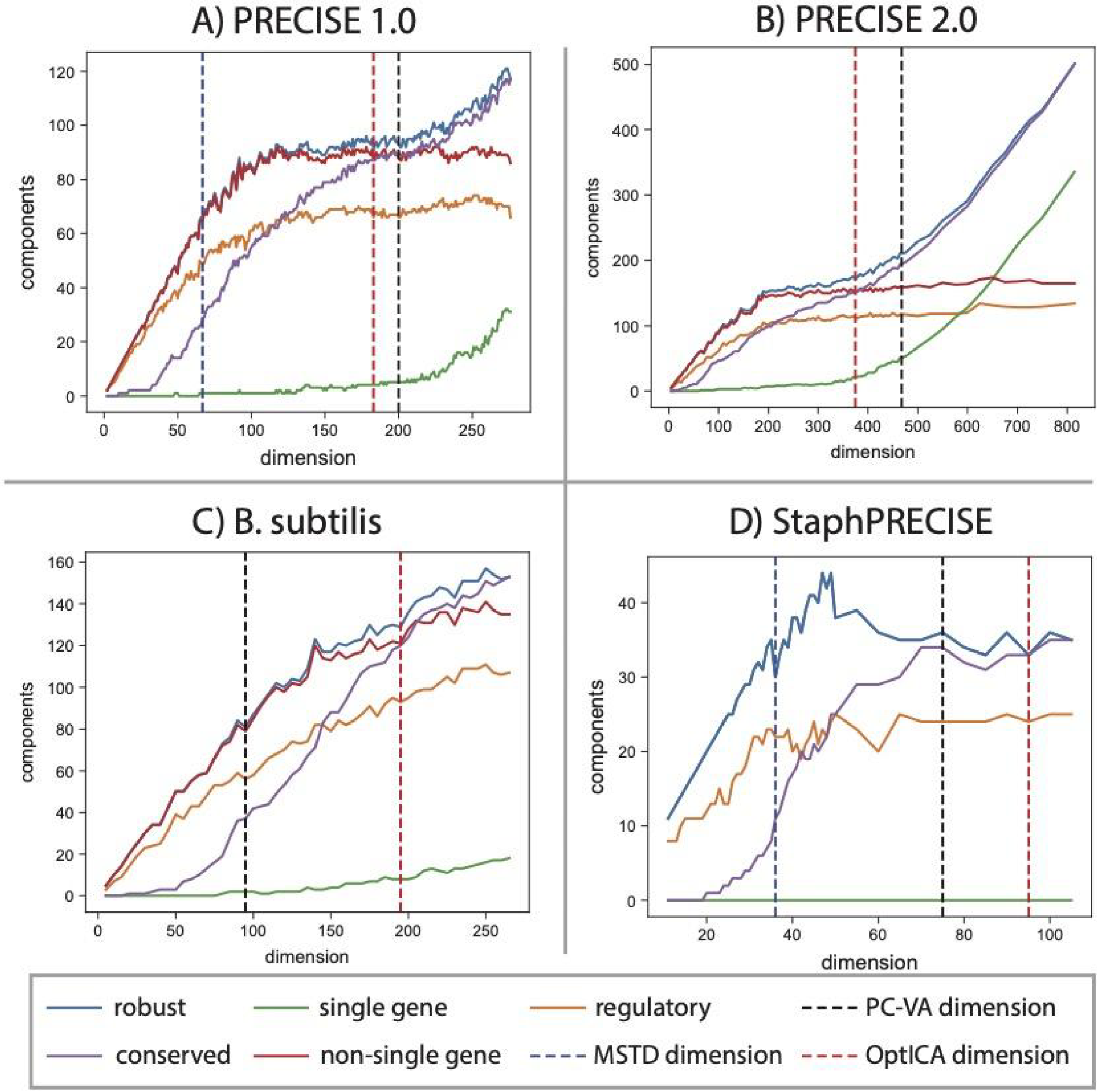
Classification of independent components across dimensions for each dataset. The PC-VA, OptICA, and MSTD dimensions are shown where applicable. The four datasets are **(A)** PRECISE 1.0 **(B)** PRECISE 2.0 **(C)** *B. subtilis*, and **(D)** *Staph*PRECISE.

Previously published ICA decompositions of three of the datasets (PRECISE 1.0, *Staph*PRECISE, and *B. subtilis*) utilized the same method for establishing dimensionality. Referred to as the PC-VA method herein, this technique sets the ICA dimension based on the number of principal components which explain a certain level of variance (e.g., 95%) in the data. However, the PC-VA method selected sub-optimal dimensions for two datasets. For the *B. subtilis* dataset, the selected dimension resulted in nearly half of the possible regulatory iModulons. Selecting a higher dimension in this case would have better captured the true TRN of the organism. On the other hand, the PC-VA method resulted in 51 single-gene iModulons from the PRECISE 2.0 dataset out of 179 total iModulons (nearly 30%), indicating that the dataset was over-decomposed.

We also attempted to compute the MSTD for each dataset. However, the algorithm did not converge on a stable dimension for either the PRECISE 2.0 dataset or the *B. subtilis* dataset. The MSTD identified for the PRECISE 1.0 and *Staph*PRECISE datasets seemed to under-decompose the datasets, as the number of robust iModulons, non-single gene iModulons, and regulatory iModulons were still rising at this dimensionality (**Figure 2a,d**).

### OptICA, a novel ICA dimensionality selection technique, controls over- and under-decomposition

Based on the observation that components are conserved across dimensions, we proposed a new method to identify the optimal dimension of the dataset. An informative decomposition would maximize the discovery of these conserved components, while minimizing the number of components with single genes. Therefore, OptICA selects the dimension at which the number of conserved components equals the number of non-single gene components.

We used three criteria to evaluate the performance of the MSTD, PC-VA dimension and the OptICA dimension: (1) the number of iModulons enriched with a transcriptional regulator (i.e., regulatory iModulons); (2) the number of single-gene iModulons; and (3) the F1-score of the regulatory iModulons (**Figure 3**). The F1-score is the harmonic average of precision and recall, which measure the number of false positives and false negatives, respectively, in iModulons as compared to published regulons in the literature. Over-decomposition would result in a high number of single-gene iModulons, whereas under-decomposition would result in a low number of regulatory iModulons and/or a low average F1-score.

**Figure 3.**
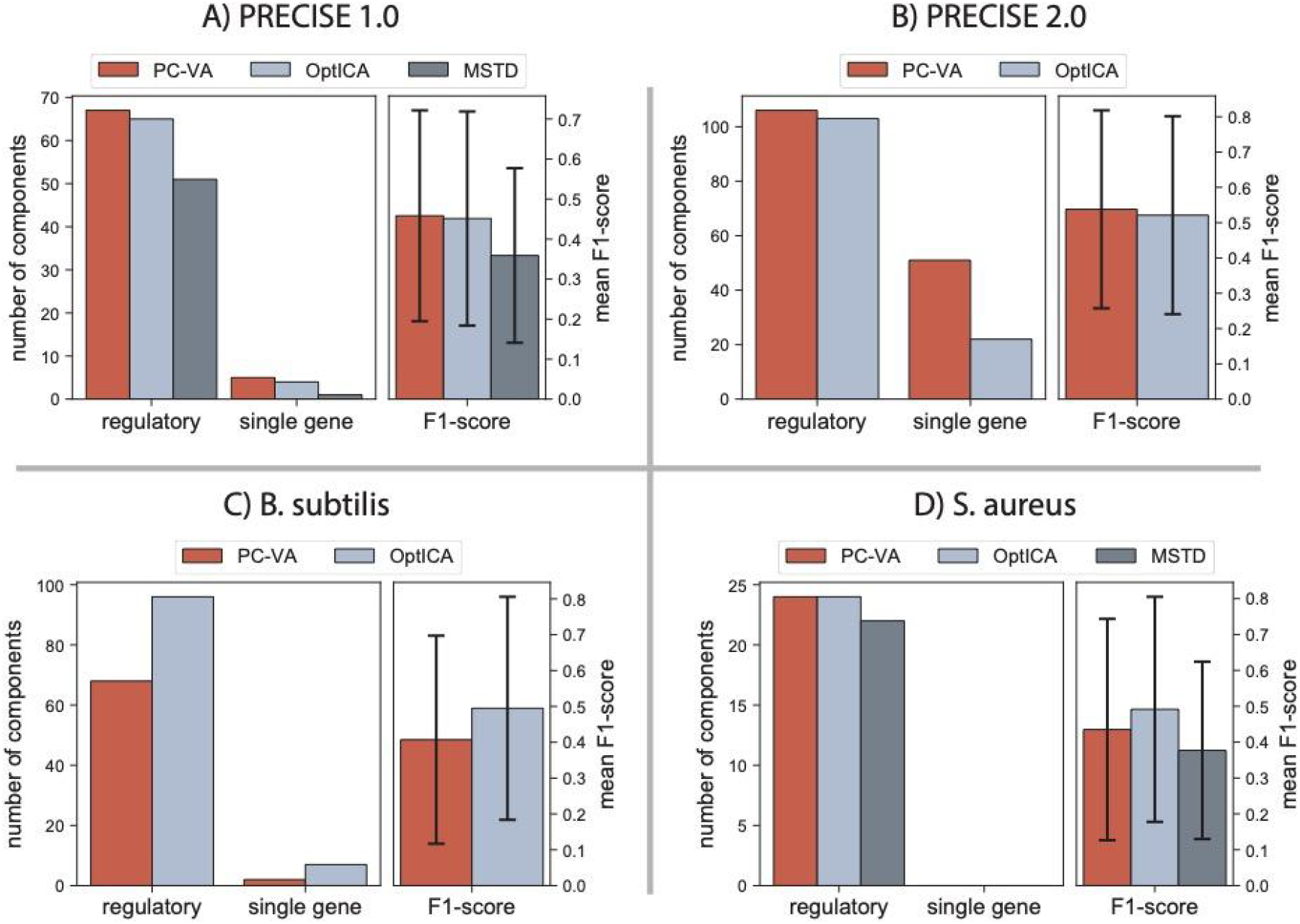
Bar charts indicating three parameters used to compare decompositions conducted at different dimensions (PC-VA, OptICA, and MSTD, when available). The left bar chart shows the number of regulatory and single gene components computed at each dimensionality, whereas the right bar chart shows the mean F1-score. Error bars represent standard deviations. Data was plotted for **(A)** PRECISE 1.0, **(B)** PRECISE 2.0, **(C)** the *B. subtilis* dataset, and **(D)** *Staph*PRECISE.

Across datasets, the OptICA method resulted in the most consistent results, selecting fully decomposed dimensions prior to the occurrence of rampant over decomposition. The PC-VA and OptICA dimensions resulted in similar decompositions for PRECISE 1.0, while the MSTD occurred at a point that captured fewer regulatory components with less congruence with associated regulons (i.e., lower F1-score) (**Figure 3a**). The OptICA dimension of PRECISE 2.0 resulted in a substantial reduction in the number of single gene components compared to the PC-VA dimension, with a minimal reduction in the number of regulatory components and mean F1-score (**Figure 3b**). Alternatively, the OptICA dimension of the *B. subtilis* dataset resulted in capturing substantially more regulatory components than PC-VA, which better modeled underlying regulatory mechanisms (i.e., higher F1-score) with a slight increase in single gene components computed (**Figure 3c**). The OptICA and PC-VA dimensions captured the same number of regulatory components in *Staph*PRECISE; however, the OptICA dimension resulted in slightly higher congruence between the regulatory components and their associated regulons, evidenced by a higher mean F1-score (**Figure 3d**).

From this analysis, we found that the OptICA dimension controlled for both over- and under-decomposition better than the PC-VA dimension and MSTD across the four datasets.

### OptICA results in more accurate TRN representations

To gain a deeper understanding of how different dimensions affect how well iModulons mirror the known TRN, we tracked the average F1-score across all dimensions for the four datasets (**Figure 4a,b**). The F1-scores seem to initially rapidly increase, and then stabilize, similar to the trajectory of the number of regulatory iModulons. Overall, the average F1-scores did not significantly differ between the PC-VA dimensions and the OptICA dimensions. This was likely because the F1-scores had neared their maximum values.

**Figure 4.**
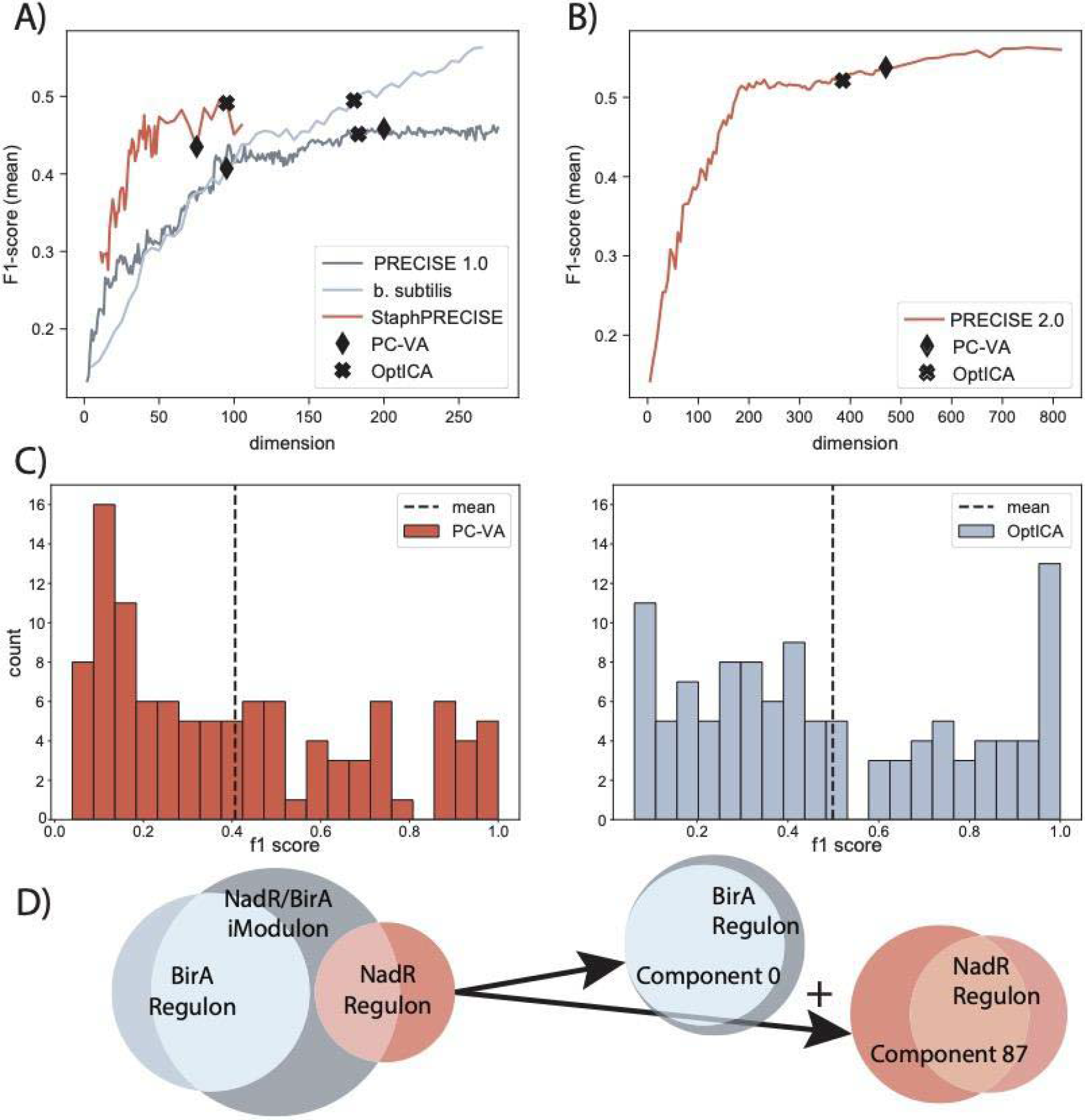
Comparing regulatory components to their associated regulon suggests the dimension selected by OptICA more accurately models the TRN. **(A)** The average F1-scores for PRECISE 1.0, *Staph*PRECISE, and the *B. subtilis* dataset at each dimensionality. **(B)** The average F1-scores for PRECISE 2.0, *Staph*PRECISE, and the *B. subtilis* dataset at each dimensionality. **(C)** The higher dimension selected by this new method applied to the *B. subtilis* dataset resulted in higher mean F1-scores and a substantial increase in components with perfect scores (exact precision and recall between the component and associated regulon). **(D)** OptICA improved F1-scores and resolved under decomposition by splitting merged components computed at the PC-VA dimension. For example, the originally published decomposition reported the NadR/BirA iModulon which contained genes belonging to both the NadR and BirA regulons. The dimension selected by this new method computed separate components which more accurately model the underlying regulatory mechanisms.

However, the OptICA dimension resulted in a meaningful increase in average F1-scores for the *B. subtilis* dataset, and a substantial increase in the number of iModulons which were perfectly aligned with a known regulon (F1-score=1.0) (**Figure 4c,d**). This improvement partially resulted from the resolution of merged components present at the PC-VA dimension (**Figure 4e**). Several iModulons from the originally published decomposition contained gene sets known to be regulated by different mechanisms; these components were effectively split at the OptICA dimension. For example, the original NadR/BirA iModulon was split into two components at the OptICA dimension which were highly congruent with their associated regulon.

## Discussion

Two important factors strongly influence the output of an ICA decomposition—the dataset of interest and the user-defined number of components to compute. Several methods have been suggested to optimally set this value in a parameter-free manner, including the MSTD and PC-VA methods described above. These methods were tested on several transcriptomic datasets and, in some cases, were found to select dimensions which under- or over-decompose the datasets evaluated, necessitating an alternative method for setting the dimensionality of ICA.

The results presented herein reveal several insights to more optimally select this specification for transcriptomic datasets. ICA was conducted on four transcriptomic datasets across a range of dimensions, showing that the overall structure of the decomposition evolves as more components are computed. In other words, as the dimensionality is increased new robust components are revealed; additionally, once a component is revealed at lower dimensions, it is well conserved across higher dimensions. This realization essentially sets a lower dimension limit for an informative decomposition which should reveal as many of these conserved components as possible.

Alternatively, an upper limit for an informative decomposition would minimize the chance for over-decomposition, which is signified by an increase in the proportion of single gene components computed. As shown by PRECISE 2.0, if a dataset is large enough, it is conceivable that each gene could be decomposed into its own iModulon, obfuscating the true structure of the dataset. The dimensionality selection method presented here, OptICA, achieves both by finding the point across the dimensionality range where the number of conserved components is equal to the number of non-single-gene components in that decomposition. Because components are well conserved across dimensions and single gene components are most often revealed at higher dimensions when over-decomposition has set in, the M-matrix at this point is likely to capture primarily the conserved, biologically relevant components.

This heuristic has two additional advantages. First, the algorithm allows for incremental, on-line learning, where the optimality of the decomposition can be assessed at each new dimension. This avoids the need to perform ICA at high dimensions, which is computationally expensive. Second, it does not require prior knowledge of the true transcriptional regulatory network since it does not rely on assessing regulatory iModulons. This enables the use of OptICA on transcriptional datasets for organisms with uncharacterized TRNs.

Overall, OptICA results in improved transcriptomic decompositions for both small and large RNA-seq datasets, avoiding both over- and under-decomposition. We validated OptICA against known transcriptional regulatory networks and found that it outperformed previously published algorithms for identifying the optimal dimensionality. OptICA is organism-invariant, and we foresee that it will assist in developing many models of transcriptional regulatory networks.

## Methods

### Conducting independent component analysis on gene expression data

The scikit-learn (v0.23.2) [18] implementation of FastICA [19] was executed 100 times with random seeds and a convergence tolerance of 10^−7^. The resulting independent components (ICs) were clustered using DBSCAN [20] to identify robust ICs, using an epsilon of 0.1 and minimum cluster seed size of 50. To account for identical with opposite signs, the following distance metric was used for computing the distance matrix:

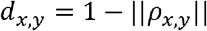

where *ρ_x,y_* is the Pearson correlation between components *x* and *y*. The final robust ICs were defined as the centroids of the cluster.

For PRECISE 1.0, which contains 278 expression profiles, the multi-start ICA process was run computing every dimension from 2 components to 276 components. For the *Staph*PRECISE dataset, every 5th dimension was analyzed from 11 components through 105 components. For the *B. subtilis* dataset, every 5th dimension was analyzed from 5 components through 265 components. For PRECISE 2.0, the process was run computing every 5th dimension from 5 components to 815 components.

### Building dimensionality trees

The cosine distance between components of each subset and those of the subsequent dimension was computed. Where this value was greater than 0.3 a connection was established between those components to build the dimensionality tree. Components from the highest dimension from each subset were similarly correlated to components of each preceding dimension. The highest of these values was used to associate a final component with each preceding component to build heat maps of conserved component occurrence in each dimension. Where the cosine distance was greater than an established threshold a final component was said to exist in a preceding decomposition. To establish these thresholds the components in the highest dimension decomposition were compared pairwise to all components computed at lower dimensions. Cosine distance was calculated for each pair and histograms of the highest values associated with a particular component in the final decomposition were plotted, resulting in a distribution of highly correlated components (**Figure S3**). The elbow point of this distribution, determined by the Kneedle algorithm [21], was used to establish a threshold correlation to classify a component as a conserved component.

### Identifying significant genes in an independent component

To perform regulator enrichments on iModulons, genes with significantly high weightings must be identified. To keep this method agnostic to the prior regulatory structure, we applied the Sci-kit learn [18] implementation of K-means clustering to the absolute values of the gene weights in each independent component. All genes in the top two clusters were deemed significant, and the set of significant genes in each independent component was called the iModulon.

### Classification of components as robust, regulatory, single gene and/or non-single gene

All components computed from a multi-start ICA decomposition, as described above, were counted as “robust components”. A component was classified as “single gene” if the highest gene weight was more than twice the next highest; the number of non-single gene components was determined by subtracting the number of single gene components from the number of robust components. The two-sided Fisher’s exact test (FDR < 10^−5^) was used to compare significant genes in each component to regulon gene sets to classify components as regulatory.

### Determining the PC-VA, MSTD and OptICA dimensions

Principal component analysis was conducted on each expression matrix, the principal components were ordered by their associated percentage of explained variance, the point at which cumulative explained variance equaled 99% determined the PC-VA dimensionality. The MSTD, or dimension at which ICA begins to compute a high proportion of unstable components, was determined as previously described [15]. The OptICA dimension was defined as the point where the number of non-single gene components was equal to the number of final components in that decomposition.

## Declarations

### Ethics approval and consent to participate

Not applicable

### Consent for publication

Not applicable

### Availability of data and materials

All data generated or analyzed during this study are available in iModulonDB (https://imodulondb.org/). Code to perform *optICA* is available at https://github.com/avsastry/modulome-workflow/tree/main/4_optICA.

### Authors’ contributions

JLM, AVS and CRL designed the study. JLM performed most of the analysis, with contributions from AVS, CRL, and SP. JLM and AVS wrote the manuscript. All authors read, edited, and approved the final manuscript.

## Acknowledgements

The authors would like to thank Kevin Rychel and Katherine Decker for helpful discussions.

## Supplementary Figures

**Supplementary Figure 1.**
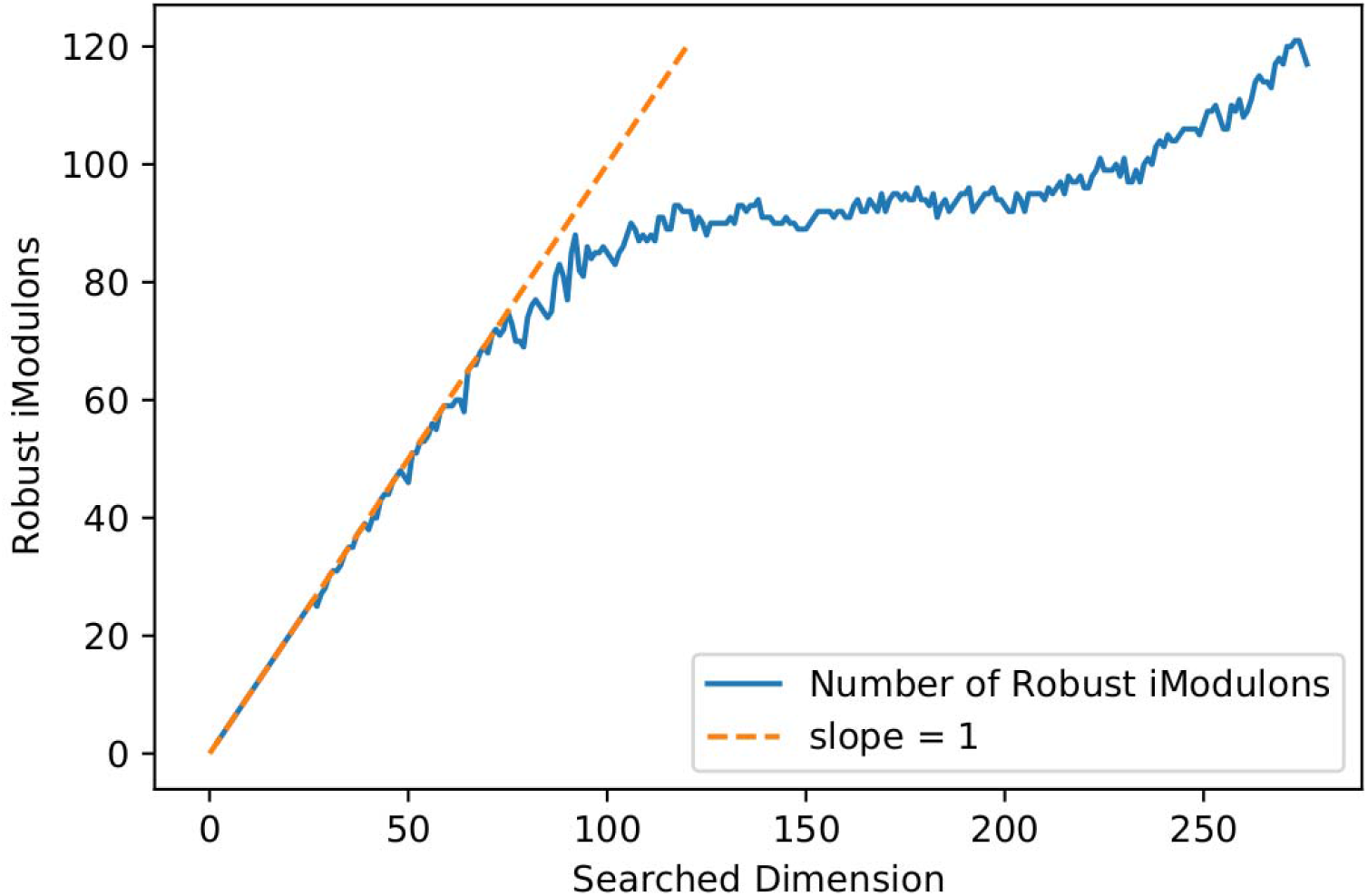
Due to density-based clustering of ICA run with randomized restarts, the number of robust components do not directly correlate with the selected dimensionality.

**Supplementary Figure 2.**
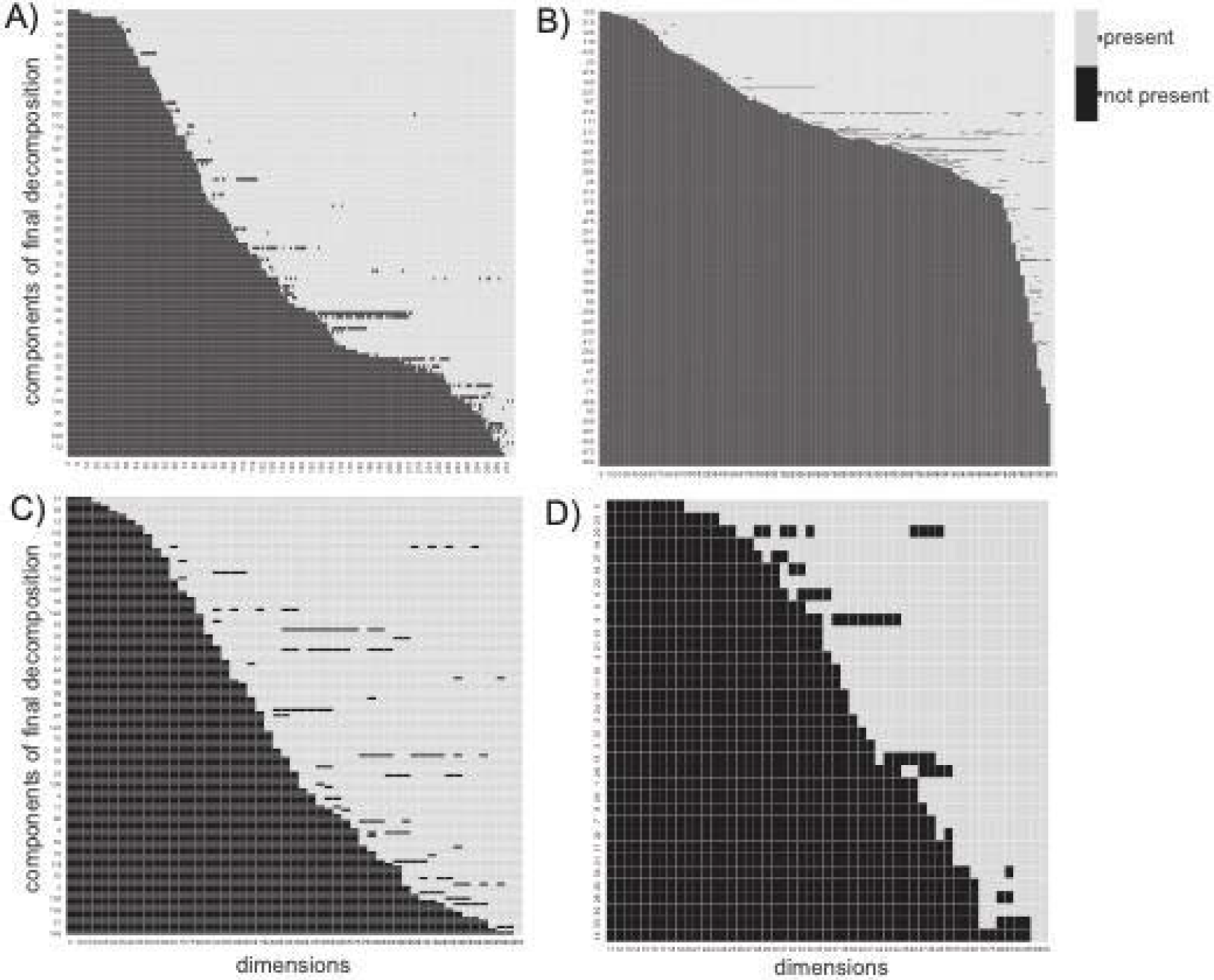
Across all dimensions components revealed in the final decomposition were found to be well conserved at the established threshold. A component in the final decomposition was said to be present in a particular decomposition where it was correlated with a component in that subset at the established threshold. Across all datasets, **(A)** PRECISE 1.0, **(B)** PRECISE 2.0, **(C)** *B. subtilis*, and **(D)** *Staph*PRECISE, components were found to be well conserved rarely dropping below their established threshold once computed at a lower dimension.

**Supplementary Figure 3.**
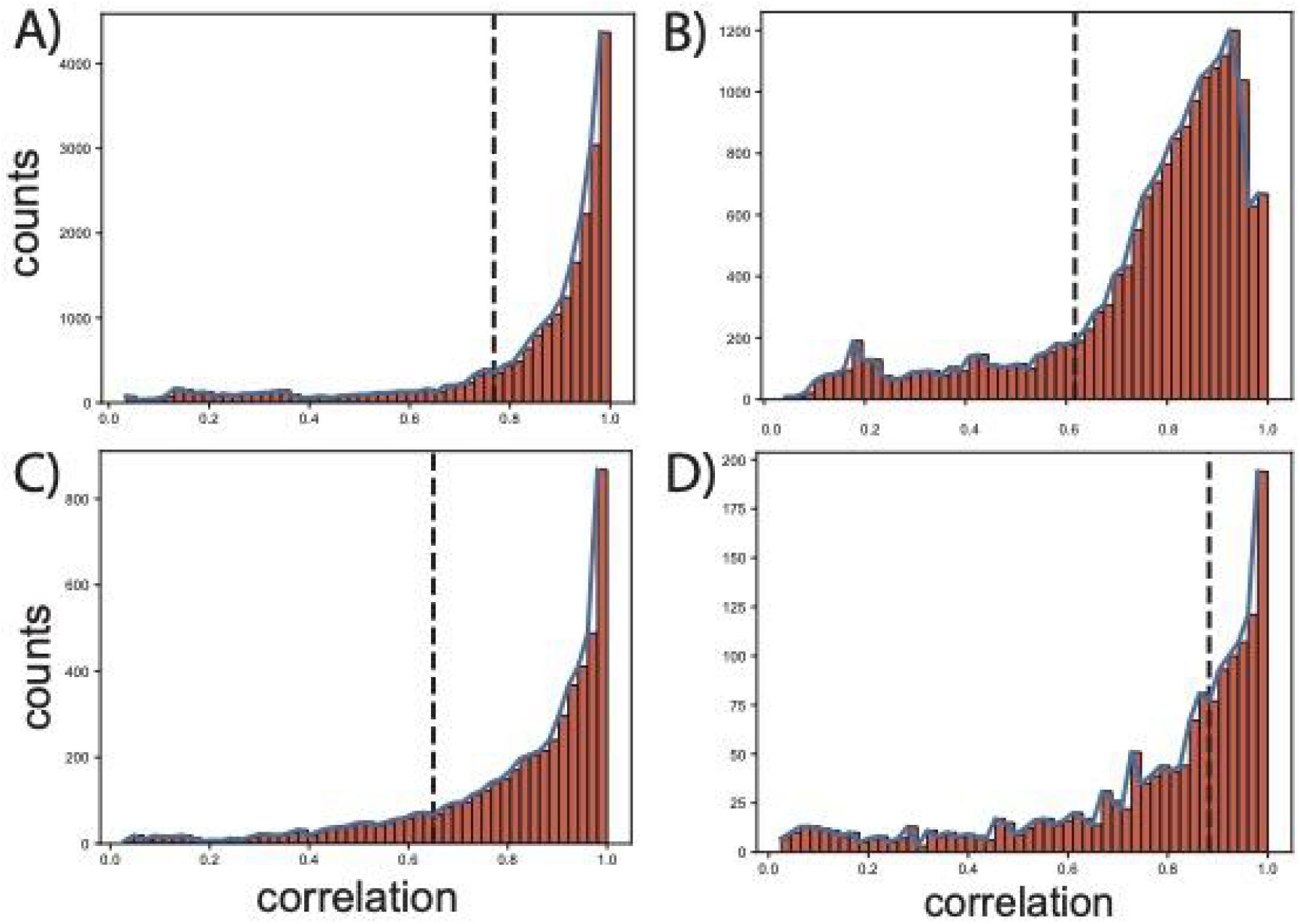
Components from the final, fully decomposed M-matrix were correlated pairwise with components of all preceding decompositions. Histograms of the highest correlations for each component across all dimensions were plotted for (A) PRECISE 1.0, (B) PRECISE 2.0, (C) *B. subtilis*, and (D) *Staph*PRECISE. The elbow point of these highly correlated values served as the threshold to classify a particular component as conserved.

